# The role of GPR87 in Pulmonary Fibrosis

**DOI:** 10.1101/2024.10.10.617569

**Authors:** Johad Khoury, Farida Ahangari, Taylor Adams, Aurelien Justet, Edward Manning, John McDonough, Thomas Barnthaler, Sabina Anderson, Fadi Nikola, Mary Lou Beermann, Jose L Gomez, Naftali Kaminski

## Abstract

**Rationale:** G- protein coupled receptor 87 (GPR87), an alternative lysophosphatidic acid (LPA) receptor previously implicated in cancer, is highly expressed in basal and aberrant basaloid cells in idiopathic pulmonary fibrosis (IPF). We sought to determine whether signaling through GPR87 is important to the development of pulmonary fibrosis.

**Methods:** Reanalysis of bulk and single cell RNA sequencing dataset was performed to confirm the increased expression of GPR87 in pulmonary fibrosis. The role of GPR87 in fibrosis *in-vivo* was assessed using global GPR87 knockout (GPR87^-/-^) and wildtype mice in the bleomycin model of pulmonary fibrosis, *in-vitro* in induced pluripotent stem cells (iPSCs) derived airway basal cells (iBC) using GPR87 siRNAs, and *ex-vivo* in human precision cut slices using disease free tissues in the fibrotic cocktail model as well as IPF tissues treated with GPR87 siRNA.

**Results:** GPR87 is highly expressed in IPF lungs, and its expression correlates with disease severity. Furthermore, It is highly expressed in basal and aberrant basaloid cells. GPR87^-/-^ mice are protected against bleomycin induced pulmonary fibrosis. GPR87 knockdown is protective against fibrosis development in normal PCLS treated with fibrotic cocktail and leads to fibrosis regression in IPF PCLS. In iBC, GPR87 knockdown leads to decreased expression of fibrosis related genes, proteins and microRNAs. GPR87 stimulation with LPA leads to the opposite results. The main downstream pathways are PI3K, mTOR, and TNF/NFkB; stimulation or inhibition of PI3K pathway mimics GPR87 stimulation or inhibition responses, respectively.

**Conclusion:** GPR87 is highly expressed in basal and aberrant basaloid cells in IPF lungs and seems to mediate profibrotic effects based on *in-vivo, ex-vivo* and *in-vitro* models of disease, suggesting that it should be studied as a potential epithelial specific therapeutic target in pulmonary fibrosis.

## Introduction

Idiopathic pulmonary fibrosis (IPF) is a chronic, fatal, progressive disease, characterized by aberrant wound healing caused by repetitive alveolar epithelial cell injury and excessive deposition of extracellular matrix proteins in the interstitial space of the lung ^1–3^. Furthermore, pulmonary fibrosis (PF) is a common final pathway of multiple primary and secondary pulmonary diseases, such as connective-tissue disease-related interstitial lung diseases, pneumoconiosis and irradiation exposure ^4^. Median survival of IPF is 3-5 years after initial diagnosis, while incidence continues to rise ^5^. Currently, there are no proven curative treatments for PF. Previously, multicenter randomized clinical trials showed promising results for two drugs, pirfenidone and nintedanib, in slowing down disease progression. However, neither drug improves lung function and, in the best case, patients are left significant pulmonary disability ^6,7^.

Growing evidence supports the role of aberrant and ectopic epithelial cells in the development of pulmonary fibrosis ^8–13^. Aberrant basaloid cells (AbBaC), first described by Adams *et al*. and Habermann *et al.*, localize to the edge of fibroblastic foci in the IPF lung ^8,9^. These cells express basal cell markers such as P63 and KRT17, but not KRT5, however; co-express known makers and regulators of fibrosis such as ITGB6, MMP7, senescence markers such as CDKN2A and CDKN2B, and epithelial–mesenchymal transition (EMT) markers such as CDH2, COL1A1, and HMGA2. Because of the location and their gene expression, these cells are thought to have a role in the development of PF ^8,9^. IPF distal airway basal cells (ABC), unlike AbBaC, carry the canonical basal cell markers, but are also strongly implicated in the pathogenesis of IPF. Cultured IPF ABC from lesions mimicking human pulmonary fibrosis in organoid models when installed into the lungs of immunosuppressed mice^11,13^. ABC signature in the transcriptome of bronchoalveolar lavage is associate with more severe outcome ^14^.

In this study we focus on G Protein-Coupled Receptor 87 (GPR87), a lysophosphatidic acid (LPA) receptor ^15,16^ expressed in ABC, and AbBaC in IPF^12,8^ and also expressed in lung, pancreas and urethral epithelial tumors ^17–19^. Rare variants of GPR87 (p.X359E and c.842–845del) were segregated in two small kindreds with familial pulmonary fibrosis, suggesting it was a disease-causing variant ^20^.

Based on GPR87’s expression in epithelial cells in human pulmonary fibrosis, it’s association with familial pulmonary fibrosis and the interest in targeting LPA and its receptors in pulmonary fibrosis ^21–24^ we studied the role of GPR87 in pulmonary fibrosis. We found that in humans, GPR87 is highly expressed in ABC or AbBaC in the IPF lung but is rarely found in the disease-free lung, and its expression is associated with more severe disease; in models of fibrosis, including *in-vitro* stimulation of epithelial cells, *ex-vivo* induction of fibrosis of human precision cut lung slices (PCLS), and *in-vivo* in mice, GPR87 was invariably highly expressed; gain of function experiments had profibrotic effects and loss of function experiments *in-vitro*, *in-vivo* and *ex-vivo* blunted fibrosis.

## Methods

### Measuring GPR87 gene expression in human lung samples

Correlation between GPR87 expression and pulmonary function tests: Gene expression data for the GPR87 gene was extracted from the publicly available Lung Genomics Research Consortium (LGRC) data at the probe-level from the Agilent-014850 Whole Human Genome Microarray 4×44K G4112F (Agilent, Santa Clara, CA), as previously described^25^. Data from individuals with idiopathic pulmonary fibrosis (IPF) and controls, including forced vital capacity (FVC) and carbon monoxide diffusion capacity (DLCO) were used for this analysis. The gene expression data is available on the Gene Expression Omnibus (GEO) database (http://www.ncbi.nlm.nih.gov/geo/) under the accession number GSE47460. Differential expression was calculated using linear mixed-effects models for each IPF group with an FDR-adjusted *P* < 0.05 considered significant for gene expression.

Correlation between GPR87 expression and surface density: Expression data was extracted from the previously published^26^ RNA-seq dataset (NCBI’s GEO GSE124685), that analyzed 95 samples, obtained from 10 IPF lungs and 6 controls. Correlation with extent of fibrosis in each sample was assessed by microCT-measured alveolar surface density (ASD) that accurately reflects tissue histology, as previously reported^26,27,28^.

### scRNAseq Data Acquisition and Processing

Raw scRNAseq data from two datasets: Adams *et al.* ^8^. and Habermann *et al*. ^9^ were downloaded from GEO (GSE136831 and GSE135893, respectively). Cutadapt was used to trim the following read 2contaminents: 5-prime terminally anchored TSO and 3-prime terminally poly(A) in the 10X 3’ v2 assayed samples from Adams *et al*.; 5-prime terminal RT-primer and 3-prime-terminal poly(T) from the 10X 5’ v1 assayed samples from Habermann *et al*. After trimming, reads were mapped using STARsolo (version 2.7.6a) to GRCh38 using GENCODE annotation release 37; cell barcode parameters were adjusted accordingly for samples from each dataset based on their respective 10X assay. Gene expression count’s from STAR’s ‘GeneFull’ output were used for downstream analysis.

### scRNAseq Analysis

Analysis was performed in R (version 4.3.3) using the package Seurat (version 4.4.0). Cells from each dataset classified as epithelial in their original analyses were isolated; data integration between the two datasets was performed with the Seurat’s reciprocal PCA (rPCA) implementation prior to UMAP embedding and cluster analysis. Cell types were collectively reclassified and a gene expression heatmap was created to demonstrate the reproducibility of the updated classifications across samples from both datasets; heatmap data was scaled independently for each dataset to avoid batch effects.

Sample-level differential expression tests between IPF and Control samples for each epithelial cell type were performed independently for each dataset. For each cell type, samples with at least 5 cells had their gene expression values averaged prior to comparison in an unpaired Wilcoxon rank-sum test. When calculating log2 fold change differences between IPF and Control, a pseudocount of 0.01 was added to each disease group’s mean to avoid distortion. Only genes with an absolute fold change greater than 0.5 were tested for significance.

### Precision cut lung slices (PCLS) single nuclei RNA sequencing

#### Sample preparation and nuclei extraction from human PCLS

We performed a time course analysis of human PCLS treated with or without Fibrotic Cocktail from day 1 (D1) through day 5 (D5). Four PCLS slices from a control donor at day 0 and at D1 to D5 stimulated with or without fibrotic cocktail were washed in cold 1X PBS and snap frozen. Nuclei were extracted using the Nuclei Isolation kit (CG000505, 10X Genomics,). Briefly and based on the manufacturer’s protocol and reagents, the tissue was dissociated on ice, centrifugated and washed. The pellet was resuspended, and cellular debris were removed. Following another centrifugation step, nuclei were resuspended and counted.

#### Single-cell barcoding, library preparation, and sequencing

Around 20,000 nuclei were loaded on a Chip G with Chromium Single Cell 3′ v3.1 gel beads and reagents (3′ GEX v3.1, 10x Genomics). Final libraries were analyzed on an Agilent Bioanalyzer High Sensitivity DNA chip for qualitative control purposes. cDNA libraries were sequenced on a HiSeq 4000 Illumina platform aiming for 150 million reads per library and a sequencing configuration of 26 base pair (bp) on read1 and 98 bp on read2.).

#### Fastq generation and read trimming

Basecalls were converted to reads with the software Cell Ranger’s (v4.0.0) implementation mkfastq. Multiple fastq files from the same library and strand were catenated to single files. Read2 files were subject to two passes of contaminant trimming with cutadapt (i) for the template switch oligo sequence (AAGCAGTGGTATCAACGCAGAGTACATGGG) anchored on the 5′ end and (ii) for poly(A) sequences on the 3′ end. Following trimming, read pairs were removed if the read2 was trimmed below 30 bp.

Paired reads were filtered if either the cell barcode or unique molecular identifier (UMI) sequence had more than 1 bp with a phred of <20. Reads were aligned with STAR (v2.7.9a) to the human genome reference GRCh38 release 99 from ensemble. Collapsed UMIs with reads that span both exonic and intronic sequences were retained as both separate and combined gene expression assays.

#### Filtering cell barcodes and quality control

After preprocessing, analysis of the *ex-vivo* human PCLS snRNA-seq data was conducted using the Seurat package (version 1.8.2). Cells with less than 750 transcripts and more that 3% mitochondrial gene ration were then removed.

#### Integration and analysis

To minimize the possible effect of potential batch correction methods, we first processed and annotated each library separately, before integrating them together and annotating them jointly. To integrate the multiple snRNA-seq datasets, we employed Robust Principal Component Analysis (RPCA). RPCA is a powerful technique for decomposing a data matrix into low-rank and sparse components. Briefly, the low-rank component represents shared biological signals across datasets, while the sparse component captures dataset-specific variations and technical noise. Based on the cellular diversity, we chose to use PCLS treated with DMSO as the reference for the integration. Following the RPCA decomposition, we utilized the low-rank component as the integrated representation of the snRNA-seq datasets. This component captured shared biological signals across conditions while mitigating dataset-specific variations. However, subsequent analyses, such as clustering and differential expression analysis, were performed on the non-integrated but normalized gene expression values. To validate the effectiveness of the integrated representation, we performed various analyses, including cell-type clustering, identification of marker genes. We also compared the results of these analyses to those obtained from individual datasets to evaluate the improvement gained through the integration process. Marker genes were computed using a Wilcoxon rank-sum test, and genes were considered marker genes if the FDR-corrected p-value was below 0.05 and the log2 fold change was above 0.5.

### *In-Vivo* Experiments

Generation of GPR87 knockout mice (GPR87^-/-^): C57BL/6N-GPR87^em1Cya^ mice were purchased from Cyagen (Santa Clara, CA) as heterozygous animals. After mating and reproducing, genotype was determined using the following primers:

Forward primer (F1): 5’-CTTCTTGTATTCCTGTGGACTG-3’
Reverse primer (R1): 5’-GGACTTCTCTTAGCCTTGCTCC-3’

Bleomycin-induced mice model of PF: Homozygous GPR87^-/-^ and wildtype littermates, age 10-12 weeks were used for experiments. Briefly, Pulmonary fibrosis was induced by intratracheal delivery of bleomycin (2 U/kg) or 0.9% saline administered by oropharyngeal instillation. Mice were euthanized and lungs were harvested on day 21 for fibrosis analysis. Mice were randomly assigned to groups. We used male mice only, 8-16 mice were included in each group, and the experiment was repeated 3 times. Although bleomycin challenge was not blinded, the results were analyzed in a blinded manner.

### Modified Ashcroft score

For Ashcroft score calculation, slides were scanned using bright field with Nikon inverted microscope at 20X magnification, at least 3 photographs per slide, followed by evaluation as described earlier with modified Ashcroft scale, by two blinded observers ^29^.

### *In-Vitro* experiments

#### Cells, cell cultures, treatment and gene knockdown

Airway basal cells (iBC) derived from induced pluripotent stem cells (iPSCs) were received from the Center for Regenerative Medicine (CReM)-Boston University ^30^. These cells were chosen because they have some similarity with both airway basal, and aberrant basaloid cells with key-signature genes: as in aberrant basaloid cells, these cells express GPR87, MMP7, ITGB6, CDH2, KRT17, P63, and their expression of KRT5 is relatively low; a unique gene signature that we could not find in other cell types. The cells were cultured into Matrigel in a density of 20X10^3^ cells/ 50 µl Matrigel per well, and exposed to a commercially available basal cell medium PneumaCult-Ex Plus, (StemCell technologies, Cambridge, MA) with supplements as previously described ^30^. iBC were exposed to different stimuli: lysophosphatidic acid 5 μM (Santa Cruz biotechnology, Dallas, TX), the Phosphoinositide 3-kinase (PI3K) inhibitor A66 32 nM (Tocris, Bristol, UK), or PI3K stimulator 740Y-P (1 mg/ml, Tocris, Bristol, UK). Cells were harvested after 10-14 days of exposure.

To perform gene knockdown, we used commercially available siRNA for GPR87 SR324210 (Origene-Rockville, MD), and for LPAR1 SR319990 (Origene Rockville, MD), and compared with scrambled RNA negative control that does not align with any published human, mouse or rat genes.

Knockdown was performed based on the manufacturer’s manual before cell seeding in Matrigel, followed by 250-500 ×10^3^ cells per well, then cells were harvested 36-72 hours after knockdown.

#### qPCR

RNA isolation, and RT-qPCR were performed as previously described ^31^. Briefly, tissues were homogenized in Qiazole using Qiagen (Hilden, Germany) mini-kit for tissue. While cells in 3D cell cultures were incubated with dispase to cleave the Matrigel, then RNA was extracted using Qiagen (Hilden, Germany) micro-kit following the manufacturer’s protocols. qPCR was done using QuantiStudio 6 Pro PCR System using TaqMan gene expression assays.

### In situ hybridization

To stain lung tissue samples, 4-μm sections of formalin-fixed, paraffin-embedded (FFPE) healthy and IPF-diseased lungs were cut with a microtome and placed on slides. These sections were stained and visualized using ACD Bio Techni-Fast Red Kit (ACD, Newark, CA). Specific probe for human GPR87 was purchased from ACD. The RNAscope assay followed the manufacturer’s instructions, and the probes were detected at a wavelength of 550 nm.

### Immunofluorescence

Immunofluorescence staining of paraffin embedded slides was done as previously described ^32^. In Brief, after rehydration, antigen retrieval was done using pH=6 retrieval buffer 95C° for 30 minutes, followed by serum blocking, and 4C° incubation with primary antibody overnight, flowed by 1 hour room temperature with secondary antibody, nuclear staining with DAPI and signal detection.

### Western Blot

Protein extraction and Western blot were performed as previously described ^31^. Briefly, cells were incubated with dispase to cleave the Matrigel, then washed 3 times with phosphate buffered saline, followed by cell lysis, protein extraction and denaturation, gel running, membrane transfer, blocking and antigen detection

### Confocal microscope

Images were captured using the Leica SP8 and Leica SP5 confocal microscopes (Leica Microsystems). Sequential imaging was conducted with a ×40 or ×63/1.4 NA objective lens. The images were analyzed with Imaris 9.7.2 software and the ImageJ bundle with Java 1.8.0.

### MicroRNA panel

MicroRNA panel was revealed using nCounter® miRNA Expression Assay Kit (Nanostring, Seattle, WA) based on the manufacturer’s protocol. The data was analyzed using nSolver 4.0 software.

### Ex-vivo Experiments

Human precision cut lung slices (hPCLS): were generated from the lungs of the IPF patients and no-disease lungs as previously described ^33^. Briefly, right-middle and lower lung lobes were inflated by injecting 2% warm (37°C) low-melting agarose, cooled down in 4°C, then tissue cores were obtained using 10 mm punch biopter, and peripheral slices (300 μm) were cut with a vibratome (Precisionary VF-300, Natick, MA).

PCLS were incubated with DMEM/F12 (Gibco), 0.1% heat deactivated fetal bovine serum (FBS) (Gibco), and incubated in 5% CO2, 37°C incubator. No disease PCLS were treated with fibrotic cocktail (FC) as previously described ^33^, consisted of 5 ng/ml recombinant transforming growth factor-β (TGF-β) R&D Systems), 5 μM platelet-derived growth factor-AB (PDGF-AB) (GIBCO), 10 ng/ml tumor necrosis factor-α (TNF-α) (R&D Systems), and 5 μM lysophosphatidic acid (LPA) (Santa Cruz Biotechnology).

At day 3, PCLS were treated with GPR87 or scrambled RNA for 24 hours, nintedanib 1 μM or vehicle (DMSO) for 3 days. PCLS were harvested at the end of day 5.

### Trichrome quantification

Microscopic scanning of the slides was conducted in a bright field using a Nikon inverted microscope at 20X magnification. Four representative images were acquired for each sample and at least 20 different random fields of view were used for collagen quantification. Trichrome quantification was done using ImageJ software with deconvoluter 2.0 plugin.

### Collagen concentration in medium

PCLS medium was collected on the end of day 5, 48 hours after the last medium change. Collagen concentration was measured using R&D systems Human Pro-Collagen I alpha 1 DuoSet ELISA kit (R&D Minneapolis, MN), following the manufacturer’s manual.

### Statistical analysis

Statistical analysis of *in-vitro*, *in-vivo*, and *ex-vivo* results was carried out in GraphPad Prism version 10. Mann– Whitney U test was used in case comparing two groups are not normally distribution, and Student’s t-test for normal distribution. For parametric set of data characterized by normal distribution, differences between two groups were assessed through unpaired Student’s t-test. One-way ANOVA with Student–Newman–Keuls post hoc test was used for pairwise comparisons of three or more groups or more than 10 per group. Efficacy experiments *in-vivo* were designed to achieve 82% power to detect 20% difference between the groups at 0.05 level of significance, 6 animals in control group and 8 in the treated group. However, due to mortality, the observed group sizes in practice deviated. All data are expressed as mean ± standard error of the mean (SEM) taking into consideration that P < 0.05 is statistically significant, except for scRNAseq and RNAseq experiments were FDR was used to control for multiple hypothesis testing.

## Results

### GPR87 is highly expressed in IPF lungs, mainly in basal and aberrant basaloid cells, and correlates with disease severity

Comparing 160 IPF patients with 132 age match control participants revealed that GPR87 is significantly increased in IPF lung cells compared with controls (Log2 Fold change = 2.12; FDR adjusted P<0.0001) (Figure 1 A); In the overall cohort GPR87 was significantly inversely correlated with forced vital capacity (FVC), forced expiratory volume in 1^st^ second (FEV1), and carbon monoxide diffusion capacity (DLCO), but because IPF patients had significantly lower pulmonary functions (Table 1) we assessed the correlation within patients IPF. GPR87 expression was inversely correlated with FVC, (r^2^=0.083, P=0.002) and with DLCO (r^2^=0.0783, P<0.001) (Figure 1B-C). GPR87 was also correlated alveolar surface density (which inversely correlates with extent of fibrosis) (r=0.404, p=0.002) in previously published analysis of differentially affected regions within the same lung ^26^. In the same dataset, we noticed a strong correlation between GPR87 and KRT5, COL7A1, KRT17, MMP7, HMGA2, CDH2 and TP63, (r correlation coefficient 0.94, 0.9, 0.89, 0.64, 0.81, 0.64, 0.61 and 0.64, P<0.001, respectively) (Supplementary Figure 1A).

**Figure 1:**
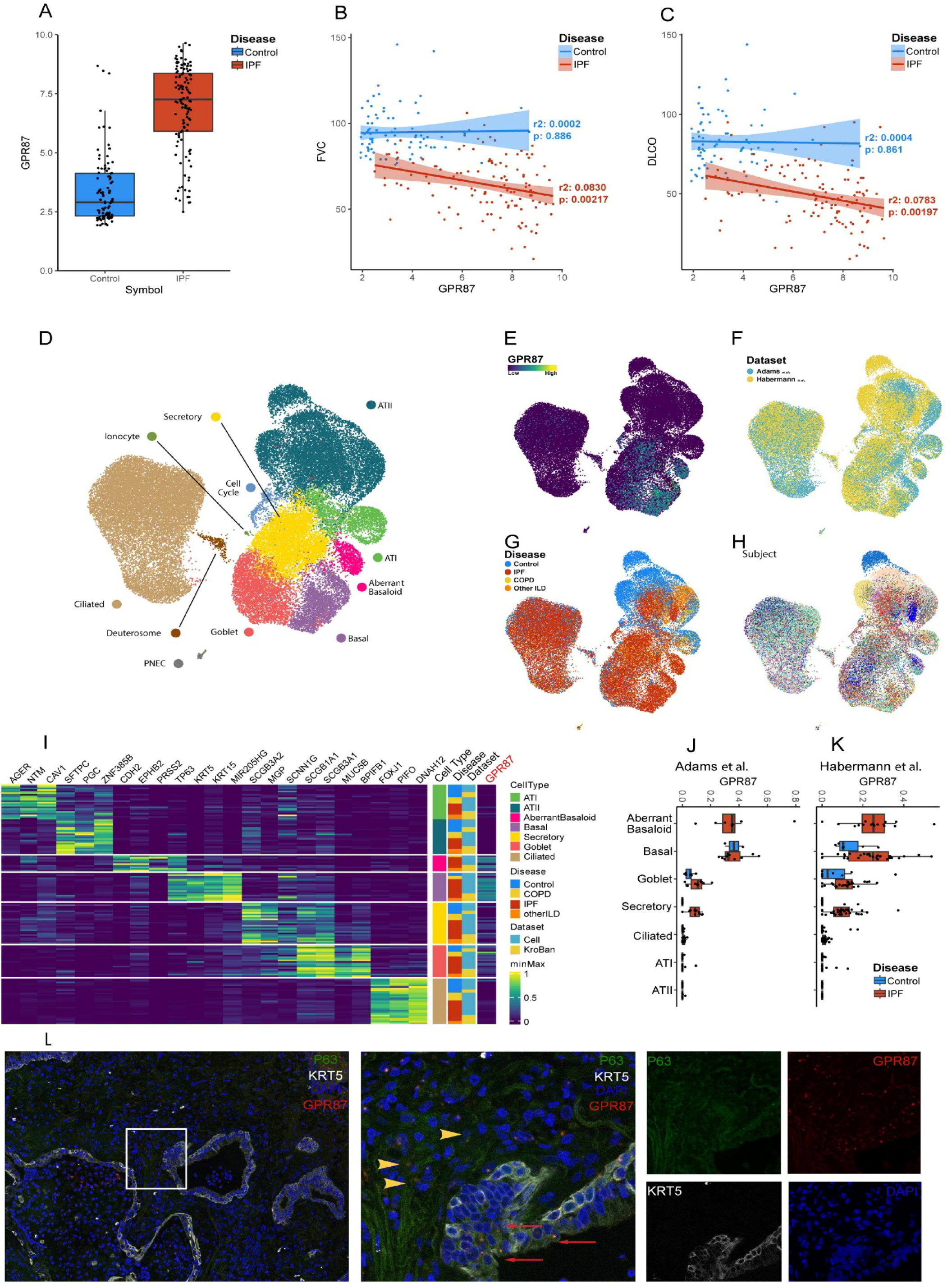
IPF: idiopathic pulmonary fibrosis; ILD: interstitial lung disease; PNEC: pulmonary neuroendocrine cells. **A**: G-coupled receptor 87 (GPR87) expression in IPF and control lungs. **B**: Correlation between GPR87 expression and FVC (Forced vital capacity). **C**: Correlation between GPR87 expression and DLCO (diffusion capacity of carbon monoxide). **D**: Uniform Manifold Approximation and Projection (UMAP) representation 47,296 epithelial cells from 107 different lungs in two different datasets. each dot represents a single cell, and cells are labeled as one of 11 discrete cell varieties. AT: alveolar type; PNEC: pulmonary neuroendocrine cell. **E**: Umap presenting GPR87 in different cells. **F**: Umap presenting distribution of cells among datasets. **G**: Umap presenting distribution of cells among lung diseases. **H**: Umap presenting distribution of cells among subject donors. **I**: Heat map of unity-normalized gene expression of curated markers observed to different cell types; each column is representative of the average expression value per cell type for one subject. **J-K** Expression of GPR87 among different cell types in Adams et al. and Habermann et al. datasets, respectively. L: IPF tissue, GPR87 in situ hybridization (Red), P63 (Green), and KRT5 (Silver) immunofluorescence and DAPI. Yellow arrow-heads showing GPR87 and P63 positive, KRT5 negative cells, probably aberrant basaloid cells, and red arrows showing GPR87, P63 and KRT5 positive cells, probably airway basal cells.

**Table 1:**
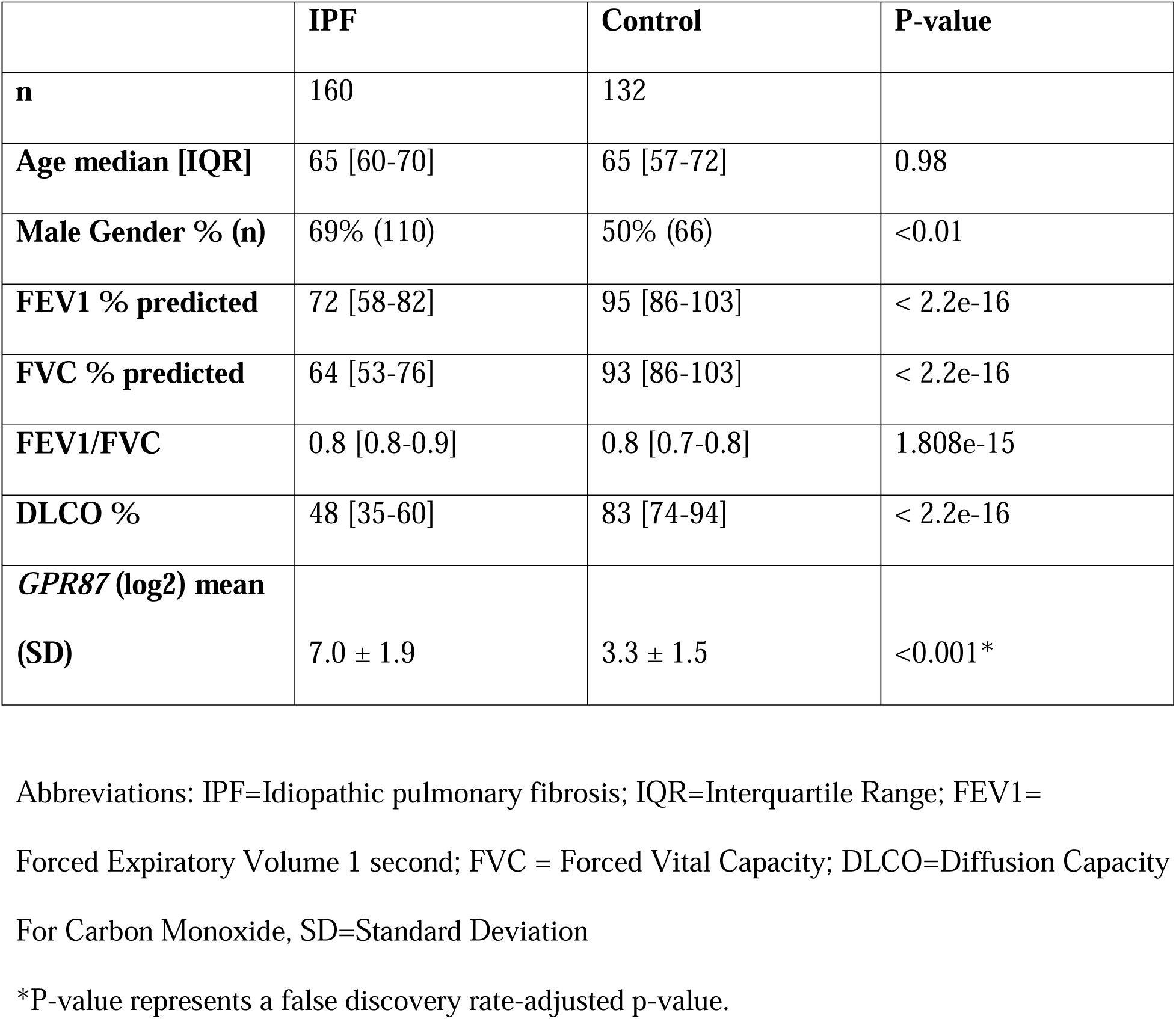
Characterization of IPF Vs Control lung samples.

|  | **IPF** | **Control** | **P-value** |
| --- | --- | --- | --- |
| **n** | 160 | 132 |  |
| **Age median [IQR]** | 65 [60-70] | 65 [57-72] | 0.98 |
| **Male Gender % (n)** | 69% (110) | 50% (66) | <0.01 |
| **FEV1 % predicted** | 72 [58-82] | 95 [86-103] | < 2.2e-16 |
| **FVC % predicted** | 64 [53-76] | 93 [86-103] | < 2.2e-16 |
| **FEV1/FVC** | 0.8 [0.8-0.9] | 0.8 [0.7-0.8] | 1.808e-15 |
| **DLCO %** | 48 [35-60] | 83 [74-94] | < 2.2e-16 |
| ***GPR87* (log2) mean**<br>**(SD)** | 7.0 ± 1.9 | 3.3 ± 1.5 | <0.001* |

Abbreviations: IPF=Idiopathic pulmonary fibrosis; IQR=Interquartile Range; FEV1= Forced Expiratory Volume 1 second; FVC = Forced Vital Capacity; DLCO=Diffusion Capacity For Carbon Monoxide, SD=Standard Deviation
\*P-value represents a false discovery rate-adjusted p-value.

47,296 epithelial cells, from 107 different lungs were reanalyzed from the Adams *et al*. ^8^ and Habermann *et al*. ^9^ single cell datasets. GPR87 is highly expressed in IPF lungs, mostly in airway basal and aberrant basaloid cells, and to a certain degree, in secretory cells (Figure 1D-K), but not in immune or mesenchymal cells. It is mostly expressed in IPF lungs in comparison with disease-free, chronic obstructive airway disease (COPD) or non-IPF interstitial lung diseases (ILD)s.

These results were validated using RNA ISH staining for GPR87 and immunofluorescence for the co-markers KRT5 and P63 (Figure 1L).

### GPR87^-/-^ mice are protected against pulmonary fibrosis

We induced fibrosis using a single dose of bleomycin (2 U/kg) administered into the lung by oropharyngeal aspiration to GPR87^-/-^ mice and wildtype littermates. On Day 21, we sacrificed the mice and harvested the lungs. The mortality of the GPR87^-/-^ mice group was lower than the WT group (25% Vs. 40%, P=0.048) (Figure 3A). All the mice in saline groups survived until the designated day. Weight loss provided additional evidence for relative protection of GPR87^-/-^ mice; the mean weight change in GPR87^-/-^ mice was significantly lower than WT mice after bleomycin (−0.8gr, −2.3 gr, respectively, fold change =2.87, p=0.05, n=19) (Figure 2B). Lung function tests revealed similar trends: after bleomycin, pressure-volume loops were higher in GPR87^-/-^ than WT, although saline groups were higher than both (Figure 3C). Similarly, lung static-compliance in GPR87^-/-^ mice was significantly higher compared to WT mice (0.04716 mL/cmH_2_O, 0.03291 mL/cmH_2_O, fold change=1.43, P=0.002, n=16). The modified Ashcroft score was significantly lower score in GPR87^-/-^ compared to WT mice after bleomycin treatment (2.6, 4.4 respectively, fold change = 1.7, P=0.04, n=19) reflecting less tissue fibrosis (Figure 2E). Quantification of collagen in the lung tissue using hydroxyproline assay revealed lower hydroxyproline in the GPR87^-/-^ compared to WT mice after bleomycin treatment group (0.71µg, 0.89 µg, respectively, fold change= 1.25, P=0.027, n=25) (Figure 2F). Relative expression of Col1A1 RNA confirmed the findings and significantly lower in GPR87^-/-^ mice compared to WT after bleomycin (1.17, 1.8 respectively, fold ratio =0.65, P=0.02, n=20) (figure 2G).

**Figure 2.**
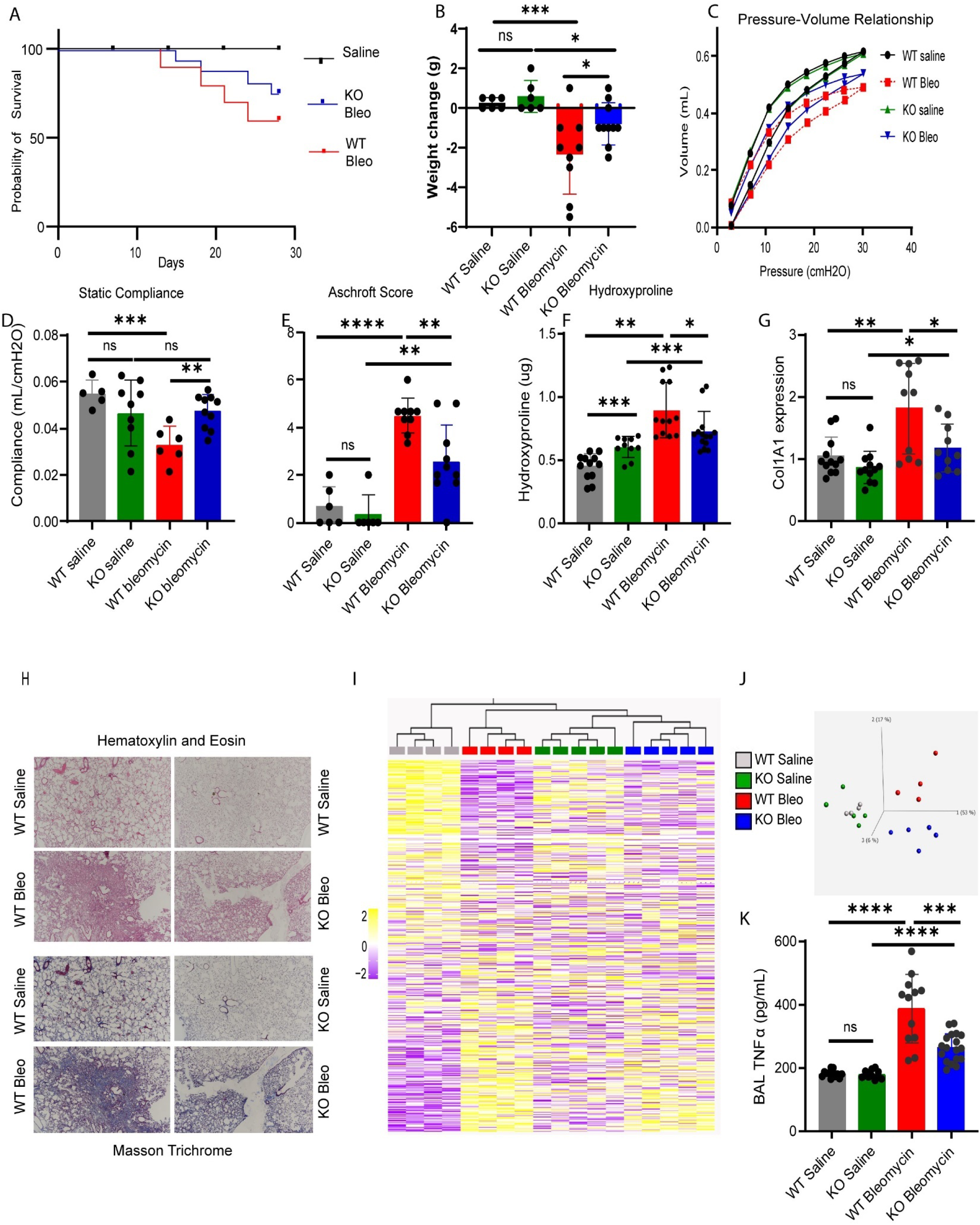
Bleo: bleomycin; WT: wildtype; KO: GPR87 global knockout; BAL: bronchoalveolar lavage; TNF: Tumor necrosis factor. **A**: Survival curve of wildtype (WT) and GPR87 KO (knockout) mice after bleomycin or saline injection. **B**: weight change between day zero and day 21 (grams) in the indicated groups of mice **C**: Pressure volume loops at day 21 in the indicated groups of mice. **D**: lung compliance in the indicated groups of mice. **E**: Modified Ashcroft score for WT and GPR87 KO mice 21 days after bleomycin or saline. **F:** Quantitative analysis of hydroxyproline in lung homogenates from indicated groups of mice. **G:** Col1A1 gene expression measured using real-time quantitative polymerase chain reaction (RT qPCR) in the indicated groups of mice. **H:** Representative images of Hematoxylin & Eosin and Trichrome staining of lung sections in the indicated groups of mice. **I:** Heatmap of all genes in the indicated groups of mice showing a similarity between WT and KO saline groups, but a difference response after bleomycin injection. **J:** three-dimensional Principal Component Analysis (PCA) plot was generated using Qlucore Omics Explorer to visualize the variance in [e.g., gene expression] data across the indicated groups of mice. **K:** TNF measured in BAL from mice lungs on sacrifice day, using ELISA.

**Figure 3.**
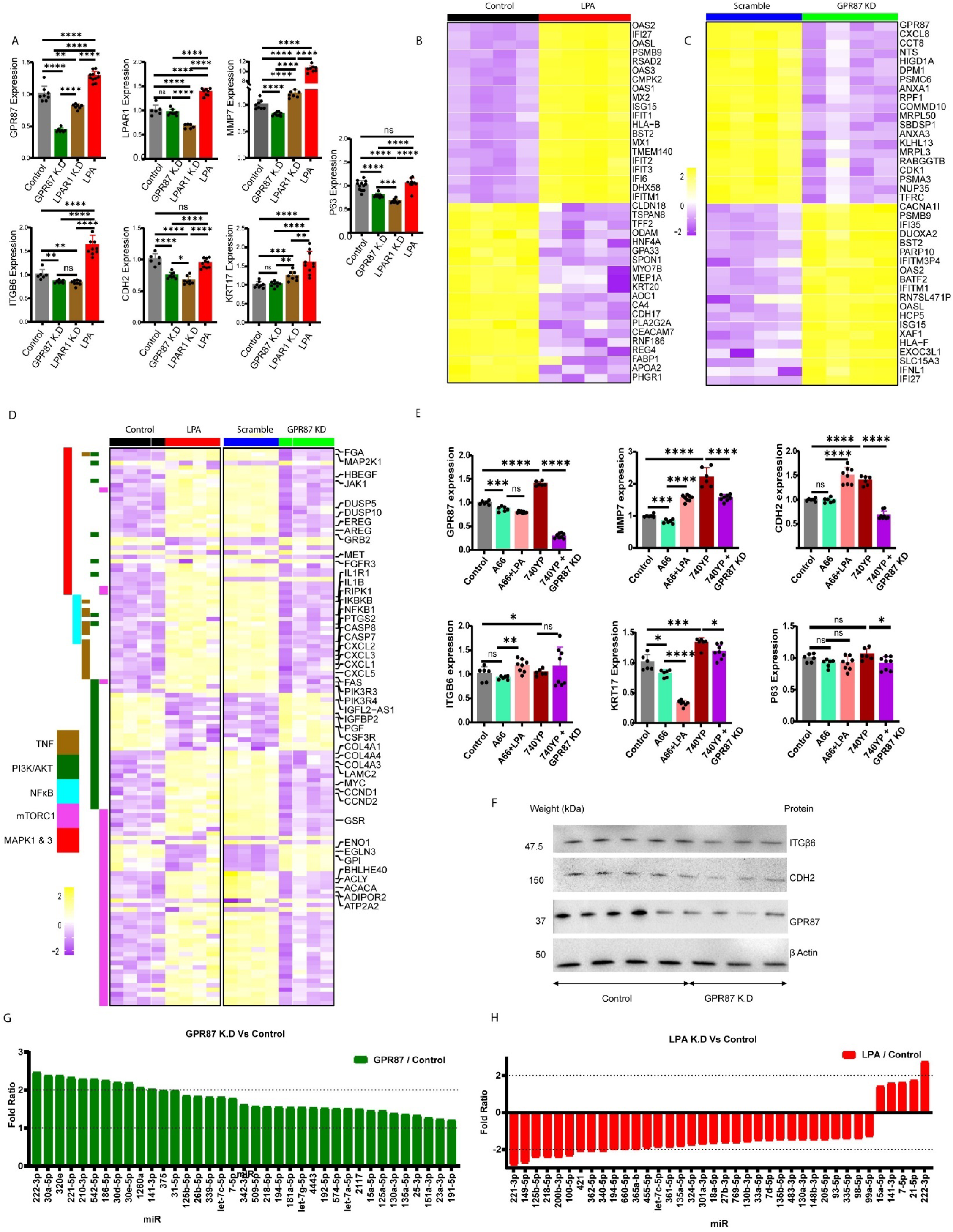
KD: knockdown (using siRNA); LPA: lysophosphatidic acid. **A:** qPCR of gene expression in induced basal cells (iBC), comparing GPR87, LPAR1, MMP7, ITGB6, CDH2, KRT17 and P63 after GPR87 knockdown, LPAR1 knockdown, lysophosphatidic acid (LPA) compared to control. **B**: Heat map of the top differentially expressing genes in iBC after LPA stimulation and GPR87 knockdown. **C-D**: Enriched pathways after GPR87 knockdown or LPA stimulation, showing involvement of PI3K, mTOR, TNF, NFkB, and MAPK1/MAPK3 pathways. E: qPCR of gene expression in iBC, after PI3K pathway inhibition using A66, with or without stimulation of LPA, and PI3K stimulation using YP-740, with or without GPR87 Knockdown. F: Western blot analysis after GPR87 knockdown in iBC, showing the expression of ITGB6, CDH2 and GPR87 protein expression. G-H micro-RNA panel showing significantly differentially expressed micro-RNAs after GPR87 knockdown, or LPA stimulation of iBC.

### Loss of GPR87 mediated protection against pulmonary fibrosis is associated with reduced TGF**β**, TNF and PI3K pathways activation

In order to reveal the pathways by which GPR87 protects against pulmonary fibrosis development in mouse model, we performed RNA bulk sequencing, which showed similarity between saline treated GPR87^-/-^ and WT mice, but were very different among GPR87^-/-^ and WT mice after bleomycin (Figure 3 I-J). Gene enrichment was performed using National Institute of Health-DAVID Bioinformatics ^34^, and KEGG PATHWAY Database ^25,35^. The most highly enriched gene clusters were related DNA repair and well documented pathways in pulmonary fibrosis DNA replication and repair and cell cycle, and TGF beta signaling pathway, it was also noted that PI3K and TNF pathways were highly enriched (Supplement figure 2).

### In airway basal cells, LPA induces fibrosis related genes, microRNAs and proteins, while GPR87 knockdown inhibits it

GPR87 was reported in the literature as a LPA receptor ^16^. To determine the effect of GPR87 stimulation, iBC were cultured with or without LPA added to the medium, although this might also stimulate LPAR1, another LPA receptor in these cells ^36^. In order to study the effect on iBC, we chose a special gene panel, that includes genes highly increased in IPF in general, and in aberrant basaloid cells in particular: MMP7, ITGB6, CDH2 ^8,9^, besides to GPR87 as the gene of interest, LPAR1 as an alternative LPA mediator ^8,9,36^, and the basal cell markers KRT17 and P63. Using qPCR to compare gene expression in LPA treated (n=11) Vs control (n=8), we noticed that LPA induced the expression of GPR87, LPAR1, MMP7, ITGβ6, and KRT17 (1.3, 1.4, 10.5, 1.6 log2 folds, respectively, P<0.001) but not P63 or CDH2 (P>0.05). (Figure 3A). On the other hand, knockdown of GPR87 by siRNA (n=8) led to decreased expression of GPR87, MMP7, ITGβ6, CDH2 and P63 (log2 fold change 0.44, 0.8, 0.85 and 0.7 respectively, P<0.01 for ITGβ6, and P<0.001 for the reminder genes). A finding also confirmed at the protein level (Figure 3F); however, did not change LPAR1 or KRT17 expression (P>0.05) (Figure 3A). To compare the effect of GPR87 to another LPA receptor expressed in these cells, we knocked down LPAR1 using siRNA, with a comparable degree of knockdown: 63% and 47% for GPR87 and LPAR1, respectively (Figure 3A). LPAR1 knockdown (n=8) led to decreases in GPR87, LPAR1, ITGB6, CDH2 and P63 (log 2 fold change 0.8, 0.7, 0.8, 0.7 and 0.7, P<0.001), but not MMP7 and KRT17, which increased by 1.2 log2 folds each, P<0.001, suggesting that LPAR1 expression contributed to the expression of ITGB6, CDH2, but probably not to MMP7.

### GPR87 knockdown mediates antifibrotic effect through mTORC1, TNF**α**, P53 and PI3K pathways

We performed bulk RNA sequencing iBC treated with LPA vs control, and iBC subjected to GPR87 siRNA knockdown vs scrambled RNA (n= 4 in each group). We analyzed bulk RNA sequencing using Qlucore software and performed student T test with FDR adjusted P<0.05 to distinguish the differentially expressed genes. 6359 genes were differentially expressed after LPA treatment, and 9520 genes after GPR87 knockdown in iBC. Top differentially expressed genes are presented in Figure 3B-C. In order to determine the pathway through which GPR87 attenuates LPA stimulation, genes which were increased with GPR87 KD and decreased with LPA stimulation (603 genes), or decreased with GPR87 KD and increased with LPA stimulation (882 genes), were noted (Supplementary figure 3A). Gene enrichment was performed using National Institute of Health-DAVID Bioinformatics ^34^ and MSigDB Hallmark 2020 ^25,37^, we identified that among the most highly enriched pathways were mTORC1, TNFα, and P53, although PI3K pathway was also highly enriched (Figure 3D). It is noteworthy to mention that these pathways overlap in PI3K pathway ^38^.

To validate GPR87 effect through PI3K pathway, we treated iBC with A66 (selective PI3K inhibitor), with (n=6) or without LPA (n=8), and with PI3K specific stimulator 740YP (n=6), and 740YP treatment followed by GPR87 knockdown (n=8). Results are shown in detail in table 2, briefly, PI3K stimulation with either LPA or 740YP led to higher expression of fibrosis related genes - GPR87, MMP7, ITGB6 and CDH2, while inhibition of the pathway using A66 or GPR87 KD, blunted this effect (Figure 3E).

**Table 2:** Gene expression in PI3K pathway stimulation and inhibition.

fold change compared to control
| Condition \ Gene | GPR87 | MMP7 | ITGβ6 | CDH2 | KRT17 | P63 |
| --- | --- | --- | --- | --- | --- | --- |
| A66 | 0.87<br>P<0.0001 | 0.83<br>P<0.0001 | 0.93<br>NS | 0.99<br>NS | 0.83<br>P=0.01 | 0.92<br>NS |
| A66 + LPA | 0.8<br>P=0.005 | 1.55<br>P<0.0001 | 1.04<br>NS | 1.5<br>P<0.001 | 1.33<br>P<0.001 | 1.06<br>NS |
| 740YP | 1.4<br>P<0.0001 | 2.2<br>P<0.0001 | 1.18<br>P=0.05 | 1.4<br>P<0.001 | 0.33<br>P<0.0001 | 0.92<br>NS |
| 740YP + GPR87<br>knockdown | 0.3<br>P<0.001 | 1.57<br>P<0.001 | 1.16<br>P>0.05 | 0.68<br>P<0.001 | 1.18<br>P=0.02 | 0.92<br>NS |

### GPR87 knockdown leads to overexpression of microRNAs related to anti-fibrotic effects, while LPA leads to an opposite result

After RNA extraction from iBC 48 hours after GPR87 siRNA knockdown, or 10 days after LPA treatment, we ran nCounter® miRNA Expression Assay Kit (Nanostring, Seattle, WA) as mentioned above. Differentially expressed microRNAs are presented in Figure 3G-H. We noticed that GPR87 knockdown led to overexpression of microRNAs related to pulmonary anti-fibrotic activity, or downregulated in IPF, including let-7 family, miR-30, mir-15, ^39^, mir-26 ^40^. On the other hand, LPA stimulation of iBC led to downregulation of Let-7 family, miR-200 family and upregulation of miR-21, which is related to pro-fibrotic activity ^41^.

### GPR87 knockdown is protective against pulmonary fibrosis in the *ex-vivo* human disease-free PCLS fibrotic cocktail treatment model

In order to study the role of GPR87 in the development of PF in human ex-vivo model, first we exposed human disease-free PCLS to fibrotic cocktail as previously described ^33^. Slices were harvested every 24 hours after the initial exposure, up to 5 days and snap frozen for future analyses. Nuclei were extracted, and a single cell RNA sequencing was performed as mentioned in the methods above. No expression of GPR87 was seen before the exposure to fibrotic cocktail, and during the first 2 days of exposure, however; it was increased starting from day 3, through the end of the experiment at day 5 cells. GPR87 was expressed in basal and aberrant basaloid, more in fibrotic cocktail (Figure 4A-E). In parallel we knocked down GPR87 in human disease-free PCLS at day 3 and harvested the tissue at day 5, as a control we used scrambled RNA, and compared to nintedanib, and to a naïve group that was not treated with fibrotic cocktail. (n=4 in each group) (Figure 4F). We quantified GPR87 RNA expression using qPCR, revealing that GPR87 knockdown group had lower expression of GPR87 in comparison to scrambled RNA (fold ratio 0.38, P<0.001). Quantification of trichrome staining (n=30 in naïve group, and 24 in other groups) showed that the trichrome stained area in GPR87 knockdown, nintedanib and naïve groups were 3.78, 4.25 and 4.23%, (statistically similar, P>0.05) but significantly lower in scrambled RNA group (7.78%, P<0.0001). (Figure 4G).

**Figure 4.**
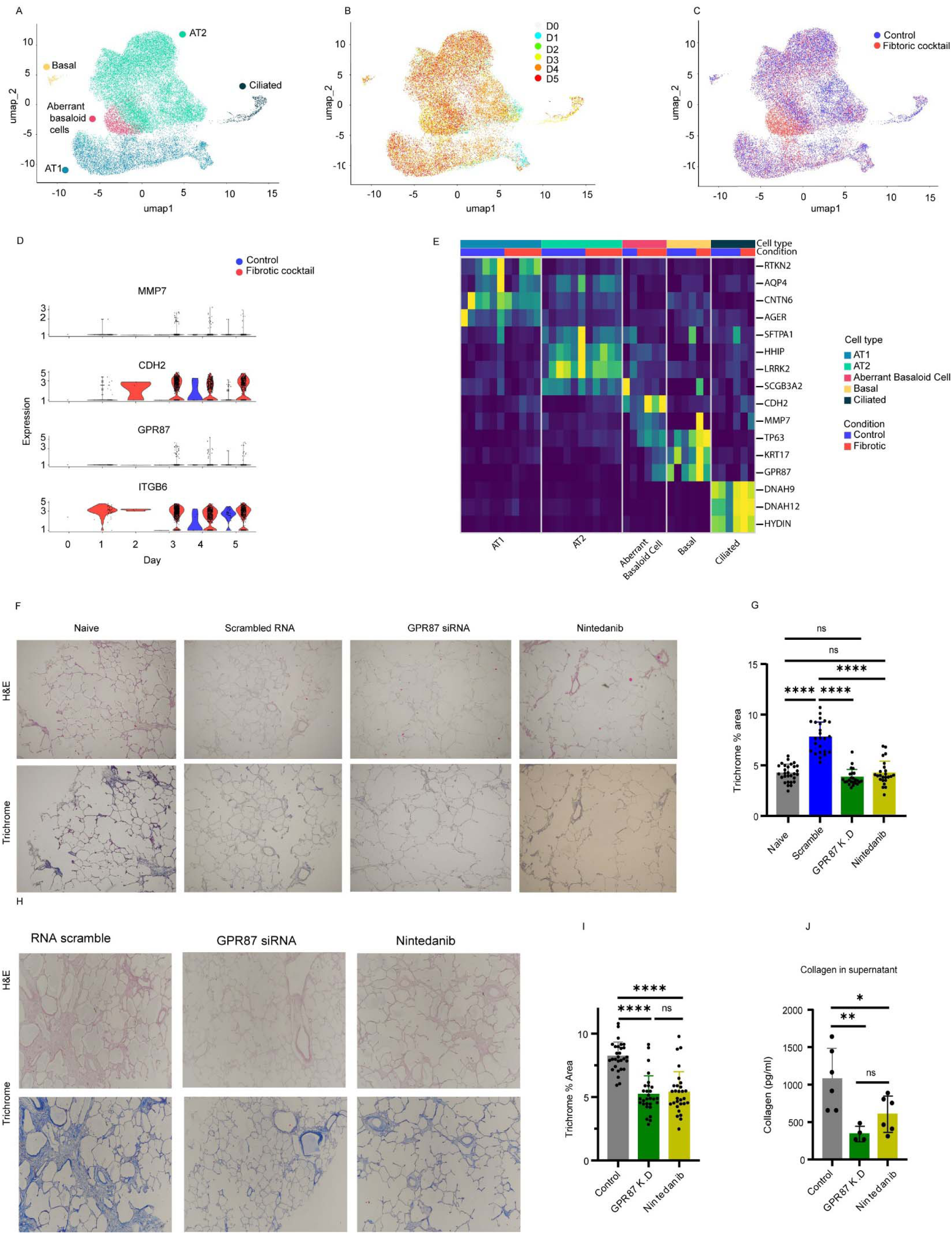
**A:** Uniform Manifold Approximation and Projection (UMAP) representation 20,000 nuclei from precision cut lung slices (PCLS), treated with fibrotic cocktail or control medium, harvested immediately after cutting, and every 24 hours up to day 5, presented: epithelial cell types, cells by day and by condition. **B**: UMAP presenting gene expression of selected genes. **C:** UMAP presenting the origin of the cells: fibrotic cocktail and control. **D:** violin plot presenting MMP7, CDH2, GPR87 and ITGB6 from day 0 through day 5. **E:** Heatmap presenting expression of some signature genes in different cell types in control and fibrotic cocktail. **F**: Hematoxylin & Eosin (H&E), and Manson trichrome stain of no-disease PCLS treated with fibrotic control medium or fibrotic cocktail with either scrambled RNA, GPR87 siRNA or nintedanib. **G**: Trichrome quantification of the previous (F) panel. **H**: Hematoxylin & eosin and Manson trichrome staining of idiopathic pulmonary fibrosis (IPF) PCLS treated with scrambled RNA, GPR87 siRNA or nintedanib. **I**: Trichrome quantification of the previous panel (F). **J**: collagen in the medium of the IPF PCLS at day 5, 48 hours after the last media change.

### GPR87 knockdown reverted fibrosis in IPF PCLS *ex-vivo* model

To determine whether GPR87 can affect pre-exisitng human pulmonary fibrosis, we knocked down GPR87 using siRNA in PCLS derived from IPF lungs, and compared to nintedanib (n=4 in each group) (Figure 4H). We quantified GPR87 RNA expression using qPCR, revealing that GPR87 knockdown group had lower expression of GPR87 in comparison to scrambled RNA (fold ratio 0.63, P<0.01). Trichrome quantification (n= 29 in each group) showed significantly lower trichrome in GPR87 knockdown and nintedanib groups than scrambled RNA group (5.2%, 5.35%, and 8.2% respectively, P<0.0001) (Figure 4I). Furthermore, measurement of collagen in the supernatant of the PCLS at the harvesting day, shows that GPR87 knockdown group (n=4) and nintedanib (n=6) had lower collagen than control (346, 606 pg/ and 1082 pg/ml, (fold ratio 0.32, 0.56) respectively, p= 0.008 and 0.03, respectively) (Figure 4J).

## Discussion

IPF airway basal cells and aberrant basaloid cells are increasingly recognized as key features and potential modulators in pulmonary fibrosis ^13,42^. Here, we demonstrate that that GPR87, an LPA receptor is highly expressed in both cell populations in IPF, while its expression in healthy cells and tissues is substantially lower. Moreover, its expression correlates with disease severity in two independent datasets. We demonstrated that the expression of GPR87 in healthy cells and murine or human tissues, is very low, but can be increased by profibrotic stimuli, such as bleomycin in mice or the profibrotic cocktail in PCLS, as well as LPA stimulation. *In-vitro,* GPR87 knockdown in iBC leads to decreased expression of fibrosis related genes, proteins and microRNAs, and GPR87 stimulation with LPA leads to a dependent increase in these genes, and microRNAs. *In-vivo* genetic knockout of GPR87 in mice is protective against development of bleomycin induced fibrosis, and *ex-vivo*, siRNA driven knockdown of GPR87 is blunts fibrosis in PCLS treated with fibrotic cocktail and IPF PCLS. Taken together our results establish GPR87 as a novel therapeutic target for pulmonary fibrosis.

In this paper, we provide the first demonstration that GPR87 has a crucial mechanistic role in the development of IPF. Although it has already been reported as an LPA receptor^15,16^, and despite the well-established involvement of LPA in the development of IPF ^23,24,43^, it was not until very recently that GPR87 was reported in IPF, specifically in airway basal cells^12^. This has been made possible with the application of cutting-edge technologies, specifically single-cell RNA sequencing that allows detection of those genes highly enriched in certain cell populations despite low overall expression in lung tissue, in this case in airway basal and aberrant basaloid cells, which growing evidence suggests their mechanistic role in IPF^13,44^. We have shown that GPR87 is a key regulator that mediates profibrotic signaling in IPF. This is seen in multiple layers: GPR87 expression correlates with disease severity, and is very rarely expressed in disease-free lungs, moreover, its expression is significantly enhanced by fibrosis stimulation, such as bleomycin, LPA or fibrotic cocktail. Importantly, GPR87 inhibition through genetic knockout, siRNA knockdown or by downstream blockade, is protective against the development of pulmonary fibrosis in *in-vitro, in-vivo* and *ex-vivo* models. Little was previously know about GPR87 expression in lung tissue, however, GPR87 was previously reported in different carcinomas ^17,19,45,46^, and correlated with cell invasion, tumor aggressiveness and metastasis in lung adenocarcinoma ^47,48^, pancreatic carcinoma ^17^ and bladder cancer ^49^. It was also described that inhibition of GPR87 using adenoviral vector, disturbed tumor proliferation in lung cancer cells ^19^; Furthermore, anti-GPR87 blocking monoclonal antibody led to regression of lung cancer in a mouse model ^45^, which highlights the mechanistic role of GPR87 in lung cancer. This leads to the conclusion that GPR87 plays a novel mechanistic role, rather than a disease marker, in the pathogenesis of IPF and presents a potential target for future therapeutic strategies.

LPA signaling in IPF has been addressed in multiple papers over the last two decades ^50,51^, however, the main focus was on LPAR1^21^ due to its high expression, especially in fibroblasts and myofibroblasts^50^. Indeed, the LPAR1 antagonist BMS-986020 showed efficacy in slowing FVC decline in IPF, however, it was associated with hepatic enzyme elevation^52^. Another phase 2 clinical trial on LPAR1 antagonist in IPF is still ongoing^53^. GPR87 and LPAR1 share some similarities, mainly conduction of LPA signaling^15,16,54^ as a mechanism for IPF development. Here we show, that GPR87 knockdown has an opposite effect to LPA treatment in basal cells. Moreover, LPA treatment leads to over-expression of GPR87, while GPR87 siRNA knockdown in disease-free PCLS, blunted the effect of fibrotic cocktail, which LPA is one of its components. Furthermore, the effect of GPR87 knockdown was similar to the effect of LPAR1 knockdown on key genes in IPF development, such as CDH2^55^ and ITGβ6^56^. However, GPR87 knockdown resulted in lower expression of MMP7, while LPAR1 knockdown did not. Moreover, GPR87 is almost exclusively expressed in IPF, specifically in airway basal and aberrant basaloid cells, whereas LPAR1 is expressed in a variety of cell types, in both healthy and IPF lungs. This might result in specificity differences while targeting GPR87 in comparison to LPAR1.

The current paradigm of IPF development points to injury to the alveolar epithelial cells as a critical event driving the development of lung fibrosis^57^. This leads to epithelial cell dysfunction and injury repair failure^58^, dysregulated epithelial-mesenchymal cross-talk^59^, abnormal activation of epithelial-mesenchymal transition (EMT) and subsequently epithelial cells lose their polarity and cell-cell adhesion properties, acquiring migratory and invasive characteristics typical of mesenchymal cells^60^. A recent study has found that IPF airway basal cells are fundamentally different from disease-free lung airway basal cells: in IPF, these cells are reprogrammed, located in areas of active remodeling and bronchiolization and adjacent to fibroblastic foci. Transcriptionally, these cells exhibit enhanced stemness, extracellular matrix sensing and epidermal growth factor signaling^13^.

Airway basal cells do migrate from airway niche and populate in fibroblastic foci and honeycomb cysts^14,61,62^. Since aberrant basaloid cells are a recently reported cell population, their origine is still not fully understood, however, they are located in highly dense fibrotic areas, present EMT and senescence markers^8,9^. GPR87 activation in these cells is highly profibrotic, mediated through well documented fibrosis pathways, particularly in the PI3K pathway, which we found to be highly enriched in both in vitro and in vivo experiments.

Furthermore, mTOR, TNF and NFκB are enriched, and overlap with PI3K pathway^38,63,64^. Based on this, it is highly likely that GPR87 activates profibrotic pathways in airway basal and aberrant basaloid cells, leading to a profibrotic effect on the cellular niche.

Our model has some limitations: first, we used one cell type, iBC, which expresses a gene signature that is similar to IPF airway basal cells and aberrant basaloid cells, since successful isolation of aberrant basaloid cells have not been reported yet. This cell model was carefully chosen to resemble the effect in IPF cells. Furthermore, the mouse model was a global GPR87 knockout, not an epithelial cell specific knockout, which may contribute to the results. To address these limitations, we used a human PCLS model, in which GPR87 expression was validated to be in basal and aberrant basaloid cells.

Taken together our results establish a novel role for GPR87 in pulmonary fibrosis, the genetic, observational and mechanistic studies we performed *in-vitro*, *in-vivo* and *ex-vivo* lead to the same conclusion, LPA signaling through GPR87 in lung epithelial cells contributes to the activation of profibrotic program, this is supported by gain and loss of function experiments in human and murine tissues. Thus, we believe that GPR87 should be considered and studied as a novel therapeutic target for IPF.

## Acknowledgments

We thank Dr. Darrell Kotton and Dr. Konstantinos Dionysios Alysandratos, Center for Regenerative Medicine (CReM), Boston University and Boston Medical Center, Boston, MA, for providing the iPSC-derived iBCs used in this study.

## References

1 Richeldi, L., Collard, H. R. & Jones, M. G. Idiopathic pulmonary fibrosis. Lancet 389, 1941–1952 (2017). 10.1016/S0140-6736(17)30866-8

2 Martinez, F. J. et al. The diagnosis of idiopathic pulmonary fibrosis: current and future approaches. Lancet Respir Med 5, 61–71 (2017). 10.1016/S2213-2600(16)30325-3

3 Lederer, D. J. & Martinez, F. J. Idiopathic Pulmonary Fibrosis. N Engl J Med 378, 1811–1823 (2018). 10.1056/NEJMra1705751

4 Wijsenbeek, M. & Cottin, V. Spectrum of Fibrotic Lung Diseases. N Engl J Med 383, 958–968 (2020). 10.1056/NEJMra2005230

5 Raghu, G. et al. Idiopathic pulmonary fibrosis in US Medicare beneficiaries aged 65 years and older: incidence, prevalence, and survival, 2001-11. Lancet Respir Med 2, 566–572 (2014). 10.1016/S2213-2600(14)70101-8

6 King, T. E., Jr., et al. A phase 3 trial of pirfenidone in patients with idiopathic pulmonary fibrosis. N Engl J Med 370, 2083–2092 (2014). 10.1056/NEJMoa1402582

7 Richeldi, L. et al. Efficacy and safety of nintedanib in idiopathic pulmonary fibrosis. N Engl J Med 370, 2071–2082 (2014). 10.1056/NEJMoa1402584

8 Adams, T. S. et al. Single-cell RNA-seq reveals ectopic and aberrant lung-resident cell populations in idiopathic pulmonary fibrosis. Sci Adv 6, eaba1983 (2020). 10.1126/sciadv.aba1983

9 Habermann, A. C. et al. Single-cell RNA sequencing reveals profibrotic roles of distinct epithelial and mesenchymal lineages in pulmonary fibrosis. Sci Adv 6, eaba1972 (2020). 10.1126/sciadv.aba1972

10 Mayr, C. H. et al. Spatial transcriptomic characterization of pathologic niches in IPF. Science Advances 10, eadl5473 (2024). doi:10.1126/sciadv.adl5473

11 Blumer, S. et al. The use of cultured human alveolar basal cells to mimic honeycomb formation in idiopathic pulmonary fibrosis. Respir Res 25, 26 (2024). 10.1186/s12931-024-02666-9

12 Heinzelmann, K. et al. Single-cell RNA sequencing identifies G-protein coupled receptor 87 as a basal cell marker expressed in distal honeycomb cysts in idiopathic pulmonary fibrosis. Eur Respir J 59 (2022). 10.1183/13993003.02373-2021

13 Jaeger, B. et al. Airway basal cells show a dedifferentiated KRT17(high)Phenotype and promote fibrosis in idiopathic pulmonary fibrosis. Nat Commun 13, 5637 (2022). 10.1038/s41467-022-33193-0

14 Prasse, A. et al. BAL Cell Gene Expression Is Indicative of Outcome and Airway Basal Cell Involvement in Idiopathic Pulmonary Fibrosis. Am J Respir Crit Care Med 199, 622–630 (2019). 10.1164/rccm.201712-2551OC

15 Ochiai, S., Furuta, D., Sugita, K., Taniura, H. & Fujita, N. GPR87 mediates lysophosphatidic acid-induced colony dispersal in A431 cells. Eur J Pharmacol 715, 15–20 (2013). 10.1016/j.ejphar.2013.06.029

16 Tabata, K., Baba, K., Shiraishi, A., Ito, M. & Fujita, N. The orphan GPCR GPR87 was deorphanized and shown to be a lysophosphatidic acid receptor. Biochem Biophys Res Commun 363, 861–866 (2007). 10.1016/j.bbrc.2007.09.063

17 Wang, L. et al. Overexpression of G protein-coupled receptor GPR87 promotes pancreatic cancer aggressiveness and activates NF-kappaB signaling pathway. Mol Cancer 16, 61 (2017). 10.1186/s12943-017-0627-6

18 Okazoe, H. et al. Expression and role of GPR87 in urothelial carcinoma of the bladder. Int J Mol Sci 14, 12367–12379 (2013). 10.3390/ijms140612367

19 Kita, Y. et al. Inhibition of Cell-surface Molecular GPR87 With GPR87-suppressing Adenoviral Vector Disturb Tumor Proliferation in Lung Cancer Cells. Anticancer Res 40, 733–741 (2020). 10.21873/anticanres.14004

20 Liu, Q. et al. The Genetic Landscape of Familial Pulmonary Fibrosis. Am J Respir Crit Care Med 207, 1345–1357 (2023). 10.1164/rccm.202204-0781OC

21 Tager, A. M. et al. The lysophosphatidic acid receptor LPA1 links pulmonary fibrosis to lung injury by mediating fibroblast recruitment and vascular leak. Nat Med 14, 45–54 (2008). 10.1038/nm1685

22 Volkmann, E. R. et al. Lysophosphatidic acid receptor 1 inhibition: a potential treatment target for pulmonary fibrosis. Eur Respir Rev 33 (2024). 10.1183/16000617.0015-2024

23 Neighbors, M. et al. Bioactive lipid lysophosphatidic acid species are associated with disease progression in idiopathic pulmonary fibrosis. J Lipid Res 64, 100375 (2023). 10.1016/j.jlr.2023.100375

24 Tanaka, T. et al. Lysophosphatidic acid, ceramide 1-phosphate and sphingosine 1-phosphate in peripheral blood of patients with idiopathic pulmonary fibrosis. J Med Invest 69, 196–203 (2022). 10.2152/jmi.69.196

25 Chen, E. et al. ENRICHR.

26 McDonough, J. E. et al. Transcriptional regulatory model of fibrosis progression in the human lung. JCI Insight 4 (2019). 10.1172/jci.insight.131597

27 De Sadeleer, L. J. et al. Lung Microenvironments and Disease Progression in Fibrotic Hypersensitivity Pneumonitis. Am J Respir Crit Care Med 205, 60–74 (2022). 10.1164/rccm.202103-0569OC

28 Tanabe, N. et al. Pathology of Idiopathic Pulmonary Fibrosis Assessed by a Combination of Microcomputed Tomography, Histology, and Immunohistochemistry. Am J Pathol 190, 2427–2435 (2020). 10.1016/j.ajpath.2020.09.001

29 Hubner, R. H. et al. Standardized quantification of pulmonary fibrosis in histological samples. Biotechniques 44, 507–511, 514-507 (2008). 10.2144/000112729

30 Hawkins, F. J. et al. Derivation of Airway Basal Stem Cells from Human Pluripotent Stem Cells. Cell Stem Cell 28, 79–95 e78 (2021). 10.1016/j.stem.2020.09.017

31 Ahangari, F. et al. microRNA-33 deficiency in macrophages enhances autophagy, improves mitochondrial homeostasis, and protects against lung fibrosis. JCI Insight 8 (2023). 10.1172/jci.insight.158100

32 Barnthaler, T. et al. Inhibiting eicosanoid degradation exerts antifibrotic effects in a pulmonary fibrosis mouse model and human tissue. J Allergy Clin Immunol 145, 818–833 e811 (2020). 10.1016/j.jaci.2019.11.032

33 Alsafadi, H. N. et al. An ex vivo model to induce early fibrosis-like changes in human precision-cut lung slices. Am J Physiol Lung Cell Mol Physiol 312, L896–L902 (2017). 10.1152/ajplung.00084.2017

34 National Institute of Health-DAVID Bioinformatics

35 KEGG.

36 Geraldo, L. H. M. et al. Role of lysophosphatidic acid and its receptors in health and disease: novel therapeutic strategies. Signal Transduct Target Ther 6, 45 (2021). 10.1038/s41392-020-00367-5

37 GSEA and MSigDB Team, <https://www.gsea-msigdb.org/gsea/msigdb/collections.jsp> (2020).

38 Peng, Y., Wang, Y., Zhou, C., Mei, W. & Zeng, C. PI3K/Akt/mTOR Pathway and Its Role in Cancer Therapeutics: Are We Making Headway? Front Oncol 12, 819128 (2022). 10.3389/fonc.2022.819128

39 Pandit, K. V., Milosevic, J. & Kaminski, N. MicroRNAs in idiopathic pulmonary fibrosis. Transl Res 157, 191–199 (2011). 10.1016/j.trsl.2011.01.012

40 Ji, X. et al. The Anti-fibrotic Effects and Mechanisms of MicroRNA-486-5p in Pulmonary Fibrosis. Sci Rep 5, 14131 (2015). 10.1038/srep14131

41 Liu, G. et al. miR-21 mediates fibrogenic activation of pulmonary fibroblasts and lung fibrosis. J Exp Med 207, 1589–1597 (2010). 10.1084/jem.20100035

42 Mayr, C. H. et al. Spatial transcriptomic characterization of pathologic niches in IPF. Sci Adv 10, eadl5473 (2024). 10.1126/sciadv.adl5473

43 Shea, B. S. & Tager, A. M. Role of the lysophospholipid mediators lysophosphatidic acid and sphingosine 1-phosphate in lung fibrosis. Proc Am Thorac Soc 9, 102–110 (2012). 10.1513/pats.201201-005AW

44 Valenzi, E. et al. Disparate Interferon Signaling and Shared Aberrant Basaloid Cells in Single-Cell Profiling of Idiopathic Pulmonary Fibrosis and Systemic Sclerosis-Associated Interstitial Lung Disease. Front Immunol 12, 595811 (2021). 10.3389/fimmu.2021.595811

45 Yasui, H. et al. Near-infrared photoimmunotherapy targeting GPR87: Development of a humanised anti-GPR87 mAb and therapeutic efficacy on a lung cancer mouse model. EBioMedicine 67, 103372 (2021). 10.1016/j.ebiom.2021.103372

46 Gugger, M. et al. GPR87 is an overexpressed G-protein coupled receptor in squamous cell carcinoma of the lung. Dis Markers 24, 41–50 (2008). 10.1155/2008/857474

47 Bai, R. et al. GPR87 promotes tumor cell invasion and mediates the immunogenomic landscape of lung adenocarcinoma. Commun Biol 5, 663 (2022). 10.1038/s42003-022-03506-6

48 Ahn, H. M., Choi, E. Y. & Kim, Y. J. GPR87 Promotes Metastasis through the AKT-eNOS-NO Axis in Lung Adenocarcinoma. Cancers (Basel) 14 (2021). 10.3390/cancers14010019

49 Zhang, X. et al. G Protein-Coupled Receptor 87 (GPR87) Promotes Cell Proliferation in Human Bladder Cancer Cells. Int J Mol Sci 16, 24319–24331 (2015). 10.3390/ijms161024319

50 Funke, M., Zhao, Z., Xu, Y., Chun, J. & Tager, A. M. The lysophosphatidic acid receptor LPA1 promotes epithelial cell apoptosis after lung injury. Am J Respir Cell Mol Biol 46, 355–364 (2012). 10.1165/rcmb.2010-0155OC

51 Oikonomou, N. et al. Pulmonary autotaxin expression contributes to the pathogenesis of pulmonary fibrosis. Am J Respir Cell Mol Biol 47, 566–574 (2012). 10.1165/rcmb.2012-0004OC

52 Palmer, S. M. et al. Randomized, Double-Blind, Placebo-Controlled, Phase 2 Trial of BMS-986020, a Lysophosphatidic Acid Receptor Antagonist for the Treatment of Idiopathic Pulmonary Fibrosis. Chest 154, 1061–1069 (2018). 10.1016/j.chest.2018.08.1058

53 Corte, T. J. et al. Phase 2 trial design of BMS-986278, a lysophosphatidic acid receptor 1 (LPA(1)) antagonist, in patients with idiopathic pulmonary fibrosis (IPF) or progressive fibrotic interstitial lung disease (PF-ILD). BMJ Open Respir Res 8 (2021). 10.1136/bmjresp-2021-001026

54 Kano, K., Arima, N., Ohgami, M. & Aoki, J. LPA and its analogs-attractive tools for elucidation of LPA biology and drug development. Curr Med Chem 15, 2122–2131 (2008). 10.2174/092986708785747562

55 Wu, S. et al. The gene expression of CALD1, CDH2, and POSTN in fibroblast are related to idiopathic pulmonary fibrosis. Front Immunol 15, 1275064 (2024). 10.3389/fimmu.2024.1275064

56 Tatler, A. L. et al. Amplification of TGFbeta Induced ITGB6 Gene Transcription May Promote Pulmonary Fibrosis. PLoS One 11, e0158047 (2016). 10.1371/journal.pone.0158047

57 Horowitz, J. C. & Thannickal, V. J. Epithelial-mesenchymal interactions in pulmonary fibrosis. Semin Respir Crit Care Med 27, 600–612 (2006). 10.1055/s-2006-957332

58 Ma, H., Wu, X., Li, Y. & Xia, Y. Research Progress in the Molecular Mechanisms, Therapeutic Targets, and Drug Development of Idiopathic Pulmonary Fibrosis. Front Pharmacol 13, 963054 (2022). 10.3389/fphar.2022.963054

59 Wisman, M. et al. Dysregulated cross-talk between alveolar epithelial cells and stromal cells in idiopathic pulmonary fibrosis reduces epithelial regenerative capacity. Front Med (Lausanne) 10, 1182368 (2023). 10.3389/fmed.2023.1182368

60 Saito, M. et al. Active mTOR in Lung Epithelium Promotes Epithelial-Mesenchymal Transition and Enhances Lung Fibrosis. Am J Respir Cell Mol Biol 62, 699–708 (2020). 10.1165/rcmb.2019-0255OC

61 Jonsdottir, H. R. et al. Basal cells of the human airways acquire mesenchymal traits in idiopathic pulmonary fibrosis and in culture. Lab Invest 95, 1418–1428 (2015). 10.1038/labinvest.2015.114

62 Xu, Y. et al. Single-cell RNA sequencing identifies diverse roles of epithelial cells in idiopathic pulmonary fibrosis. JCI Insight 1, e90558 (2016). 10.1172/jci.insight.90558

63 Gong, H. et al. Eupatilin inhibits pulmonary fibrosis by activating Sestrin2/PI3K/Akt/mTOR dependent autophagy pathway. Life Sci 334, 122218 (2023). 10.1016/j.lfs.2023.122218

64 Hou, J. et al. TNF-alpha-induced NF-kappaB activation promotes myofibroblast differentiation of LR-MSCs and exacerbates bleomycin-induced pulmonary fibrosis. J Cell Physiol 233, 2409–2419 (2018). 10.1002/jcp.26112

