## Supplementary for "The role of GPR87 in Pulmonary Fibrosis"

**Supplementary Methods:**

**Measuring GPR87 levels in human lungs and correlation with physiologic measurements:**

Gene expression data for the *GPR87* gene was measured in The Lung Genomics Research Consortium (LGRC) Cohort at the probe-level from the Agilent-014850 Whole Human Genome Microarray 4x44K G4112F (Agilent, Santa Clara, CA), used in the LGRC cohort as previously described (1). Data from individuals with idiopathic pulmonary fibrosis (IPF) and controls, including forced vital capacity (FVC) and carbon monoxide diffusion capacity (DLCO) were used for this analysis. The gene expression data is available on the GEO database (http://www.ncbi.nlm.nih.gov/geo/) under the accession number GSE47460.

**In situ hybridization (ISH):**

To stain lung tissue samples, 4-μm sections of formalin-fixed, paraffin-embedded (FFPE) healthy and IPF-diseased lungs were cut with a microtome and placed on slides. These sections were stained and visualized using ACD Bio Techni-Fast Red Kit (ACD, Newark, CA) following the manufacturer’s protocol. In brief, slides were incubated in 60^◦^ C for 60 minutes, and cooled down in room temperature overnight. The next day, after rehydration process, sections were incubated with hydrogen peroxide, followed by retrieval buffer, probe hybridization for 2 hours, amplification, and color detection. Specific probe for human GPR87 was purchased from ACD. Following color detection, sections were blocked with 2.5% goat serum, then primary antibodies overnight in 4^◦^ C, followed by the next immunofluorescence staining protocol steps as previously described (2) (immunofluorescence section). The ISH probes were detected at a wavelength of 550 nm.

**Immunocytochemistry/Immunofluorescence:**

Immunohistochemistry (IHC) was performed as previously described (2). In brief, slides were rehydrated, heat-mediated antigen retrieval was performed in citrate buffer (pH=6, 95C^o^ for 30 minutes) followed by 2.5% goat serum blocking in room temperature for 1 hour. Then samples were incubated with primary antibodies overnight at 4°C after blocking. To reduce autofluorescence, slides were incubated in TrueView reagent for 3 minutes before mounting with a mounting medium containing DAPI. IHC quantification was performed in a blinded fashion by using ImageJ’s deconvolution tool.

**Antibodies used:**

| Antigen | Manufacturer | Host | Concentration | Cat. |
| --- | --- | --- | --- | --- |
| KRT5 | NJS Bioreagents (San Diego, CA) | Mouse | 1:100 | V2176SAF |
| P63 | NJS Bioreagents  (San Diego, CA) | Rabbit | 1:50 | V3815 |

**Mice Pulmonary function tests:**

Mice were anesthetized with 18% urethane 300 µL intraperitoneal injection and paralyzed with an intraperitoneal injection of pancuronium bromide (1 mg/kg) depth of anesthesia was assessed as the lack of response to a toe pinch, with supplemental injections given as needed. Once adequate anesthesia and paralysis was achieved, the mice were placed in a supine position, and the trachea was canulated with 20G tube, through a ventral incision in the rostral-most part of the trachea and advanced 3 mm caudal to the incision. Mice were mechanically ventilated using the SCIREQ FlexiVent apparatus with 150 breaths/min, a tidal volume of 10 mL/kg body mass, and a positive end-expiratory pressure of 3 cmH_2_O prior to lung function measurements.

With the maximal vital capacity perturbation (called total lung capacity by SCIREQ), the inspiratory capacity of the lungs was determined using the SCIREQ software (Flexiware v.7.6, Service Pack 6). Forced oscillation perturbations (“quickprime-3”) subsequently measured tissue damping, reflecting energy dissipation within the lung parenchyma. Pressure-volume loops were calculated through quasi-static stepwise pressure-guided measurements of pressure *P* and volume *V*. SCIREQ software calculated static compliance by fitting the Salazar–Knowles equation (3). All maneuvers were performed till three consecutive consistent measurements were achieved per animal. A coefficient of determination of 0.9 was the lower limit for accepting a measurement.

**RNA extraction:**

In induced basal cells (iBC) were incubated with dispase 2 mg/ml for 15 minutes, then mixed by pipetting to cleave the Matrigel. The complex was centrifuged and the supernatant was discarded. Qiazole (Qiagen (Hilden, Germany)) was added, and mixed by vortexing. Using Qiagen miRNeasy micro-kit, following the manufacturer’s manual, RNA was extracted.

For tissue RNA extraction, tissue was homogenized using a D1000 Hand-Held Homogenizer (Benchmark, Tempe, AZ), followed by RNA extraction using miRNeasy mini-kit, following the manufacturer’s manual.

The purity of the RNA was verified using a NanoDrop at 260 nm, and the quality of the RNA was assessed using the Agilent 2100 Bioanalyzer (Agilent Technologies)

**Gene knockdown using siRNA:**

Target genes were knockdown before being seeded in Matrigel, following the manufacturer’s manual. The three different sequences of each gene were mixed together in the 1x transfection buffer, and siTran 2.0 siRNA transfection reagent. After incubation with transfection cocktail, cells were seeded into Matrigel, and harvested after 36-48 hours.

SiRNA GPR87 sequences:

SR324210A rArUrUrCrUrUrCrArGrUrUrGrUrGrArUrGrCrArCrUrGrUrArArGrC

SR324210B rGrUrArCrArUrArUrCrGrArUrUrCrCrArArCrArArArCrArArUrArA

SR324210C rArGrArArUrArArArCrUrUrGrArArGrUrArCrCrArArGrGrUrCrCrA

Negative control:

Negative control:Sense 5’ rCrGrUrUrArArUrCrGrCrGrUrArUrArArUrArCrGrCrGrUAT Antise 5’ rArUrArCrGrCrGrUrArUrUrArUrArCrGrCrGrArUrUrArArCrGrArC .

**qPCR:**

Real-time Quantitative Reverse Transcription-Polymerase Chain Reaction (qPCR) for RNA expression:

Relative expressions of messenger RNAs from all in vitro, in vivo, and ex vivo experiments were determined by real-time quantitative reverse transcription-polymerase chain reaction (qRTPCR) on QuantiStudio 6 Pro PCR System using TaqMan gene expression assays. Reverse transcription with random primers and subsequent PCR were performed with TaqMan RNA-CtoT one-step kit (Applied Biosystems). Raw data for cycle threshold (Ct) values were calculated using the QuantiStudio 6 Pro PCR with an automatically set baseline. The results were analyzed by the ΔΔCt method and GUS-B (β-glucuronidase) or GAPDH (Glyceraldehyde 3-phosphate dehydrogenase) were used as a housekeeping gene. Fold change was calculated by taking the average over all the control samples as the baseline. All the probes used in this study were purchased from Thermo Fisher Scientific.

**Western Blot:**

For Western blot, isolated cells were lysed and in M-Per (Thermo Fisher Scientific) added phosphatase and protease inhibitor (100 µl per 1X10^6^ cells, Abcam, ab201119) on ice for 10 minutes. Protein content was measured using ELISA and Pierce BCA Protein Assay Kit (Thermo Scientific) and denaturation was performed at 95°C for 5 min in the presence of mercaptoethanol and Laemmli buffer. 20 µg protein per lane was loaded onto a 4-20% gel (bio rad) and samples were run at 25 mA followed by transfer on PVDF membranes using the Trans-Blot Turbo Transfer System (Bio-Rad). Membranes were washed and blocked in 5% dry milk (American Bio-Inc) for 60 min followed by incubation overnight (at 4°C) with primary antibody according to the manufacturer’s instruction. Primary antibodies used as follows; β-actin (sc-47778) Santa Cruz Biotechnology (Dallas, TX), GPR87 (ab272873) Abcam (Cambridge, United Kingdom), CDH2 (V3391-) NSJ bioreagents (San Diego, CA), Signal was detected using appropriate HRP conjugated secondary antibody (1:1000 for 1h at room temperature) using ECL substrate (Bio-Rad). Visualization was performed using an enhanced chemiluminescent detection kit (Bio-Rad, Hercules, CA, USA). Quantification of blots was done by densitometry using Bio-Rad Image Lab Software 6.1 (Bio-Rad Laboratories) and actin as a loading control.

**Bulk seq gene analysis and enrichment:**

Poly-A mRNA was enriched from 200 ng of total RNA, followed by fragmentation of the mRNA to ~200-300 bp and cDNA synthesis. Equimolar amounts of indexed libraries were pooled and loaded onto an Illumina HiSeq platform for paired-end 100 bp sequencing. Data was analyzed using Qlucore omics explorer 3.8 and normalized using Trimmed Mean of M-values (TMM), considering the gene length of each gene.

In induced basal cells (iBC) Genes which have significant differential expression (P<0.05) between treatment and control groups were selected from two experiments: LPA vs Control, and GPR87 KD vs Control. Then genes which were upregulated when treated with LPA and downregulated with GPR87 KD or downregulated when treated with LPA and upregulated with GPR87 KD, were selected.

Genes where enriched using Gene enrichment was performed using National Institute of Health- DAVID Bioinformatics and Enrich MSiGDB hallmark (can be found at: [https://maayanlab.cloud/Enrichr/enrich MSigDB Hallmark 2020](https://maayanlab.cloud/Enrichr/enrich%20MSigDB%20Hallmark%202020)).

For mouse experiments, two-way anova was used to compare the groups, with P<0.05 considered significant. Furthermore, a direct comparison between knockout (KO) bleomycin and wildtype (WT) bleomycin groups was done using student T test with P<0.05 considered significant. all differentially expressed genes between these two groups were ran through Gene enrichment was performed using National Institute of Health- DAVID Bioinformatics or KEGG pathway, wikipathway selected.

**Human PCLS processing and culturing:**

Human precision cut lung slices (hPCLS) were generated from the lungs of IPF patients and no-disease lungs as previously described (4). Briefly, right-middle and lower lung lobes were inflated by injecting 2% warm (37°C) low-melting agarose, cooled down in 4°C, then tissue cores were obtained using 10 mm punch biopter, and peripheral slices (300 μm) were cut with a vibratome (Precisionary VF-300, Natick, MA).

**References:**

1.	Bauer Y, Tedrow J, de Bernard S, Birker-Robaczewska M, Gibson KF, Guardela BJ, et al. A novel genomic signature with translational significance for human idiopathic pulmonary fibrosis. Am J Respir Cell Mol Biol. 2015;52(2):217-31.

2.	Barnthaler T, Theiler A, Zabini D, Trautmann S, Stacher-Priehse E, Lanz I, et al. Inhibiting eicosanoid degradation exerts antifibrotic effects in a pulmonary fibrosis mouse model and human tissue. J Allergy Clin Immunol. 2020;145(3):818-33 e11.

3.	Salazar E, Knowles JH. An Analysis of Pressure-Volume Characteristics of the Lungs. J Appl Physiol. 1964;19:97-104.

4.	Alsafadi HN, Staab-Weijnitz CA, Lehmann M, Lindner M, Peschel B, Konigshoff M, et al. An ex vivo model to induce early fibrosis-like changes in human precision-cut lung slices. Am J Physiol Lung Cell Mol Physiol. 2017;312(6):L896-L902.

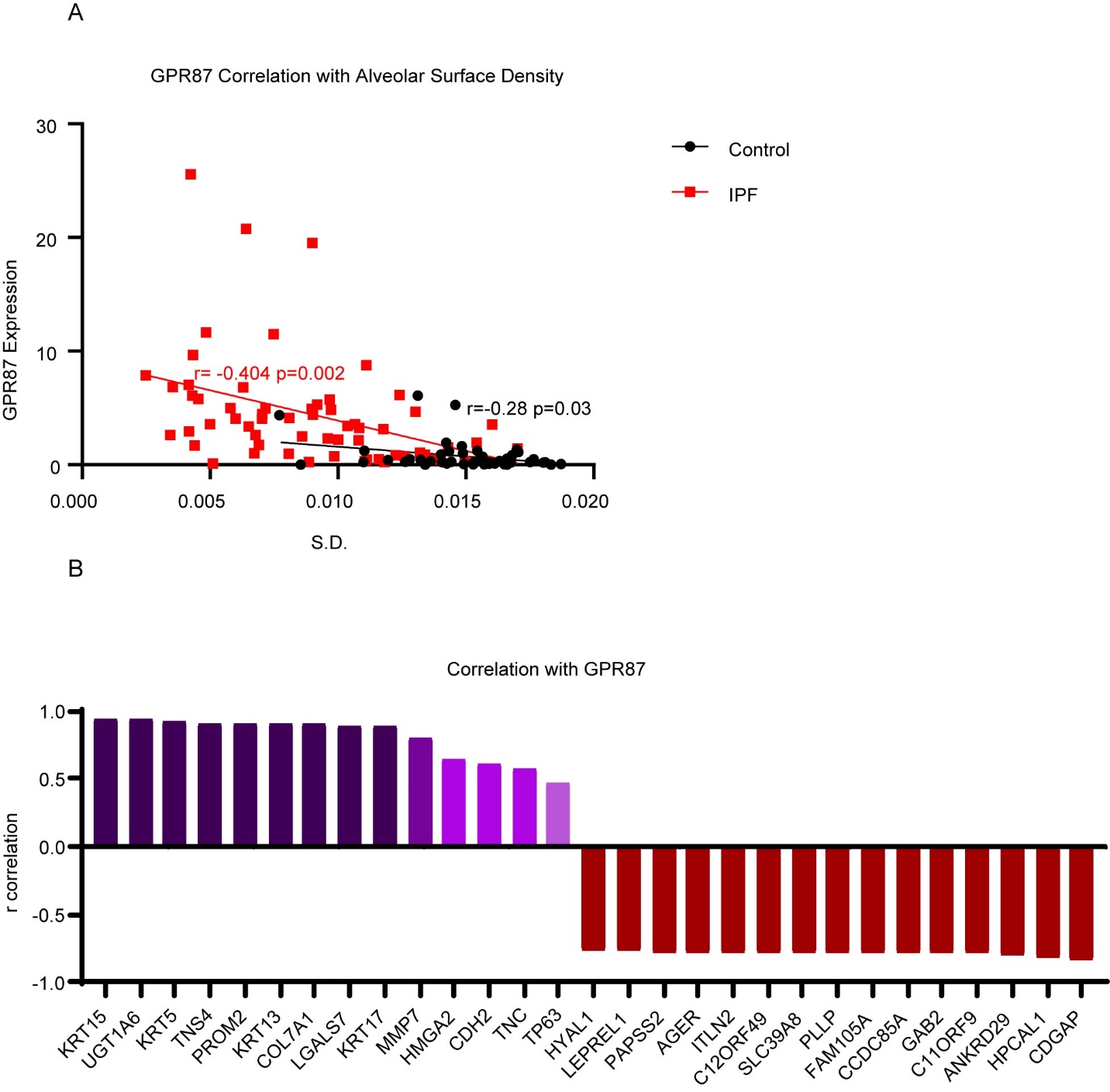
Supplementary figure 1:

A: correlation between GPR87 and alveolar surface density in LGRC dataset. B: Correlation between GPR87 expression and top directly or inversely correlated genes, and genes of interest.

Supplementary Figure 2:

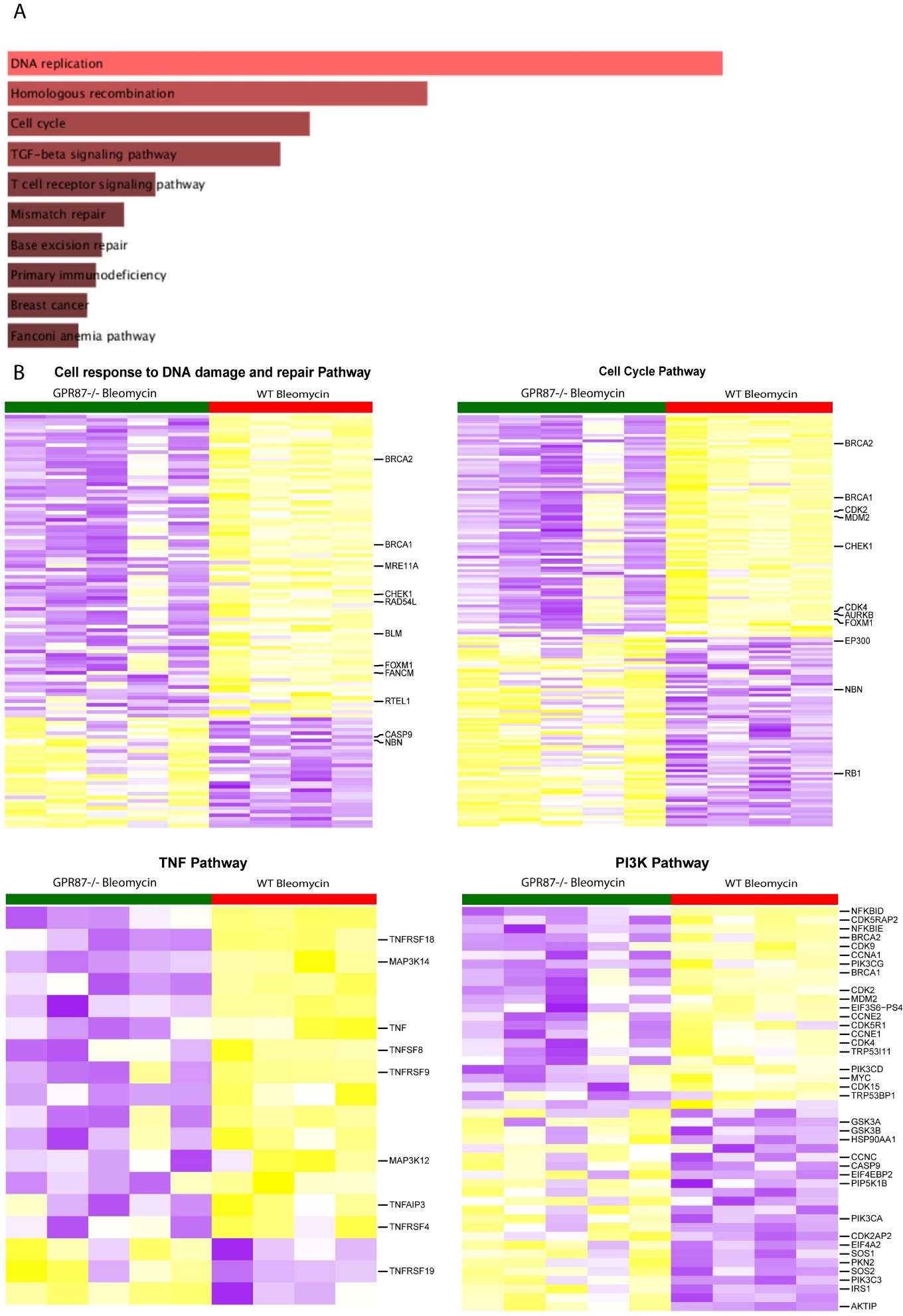

A: Top enriched pathways in bleomycin GPR87 knockout mice and wildtype mice.

B: Heatmaps of differentially expressed genes in bleomycin GPR87 knockout mice and wildtype mice in different pathways

Figure 3:

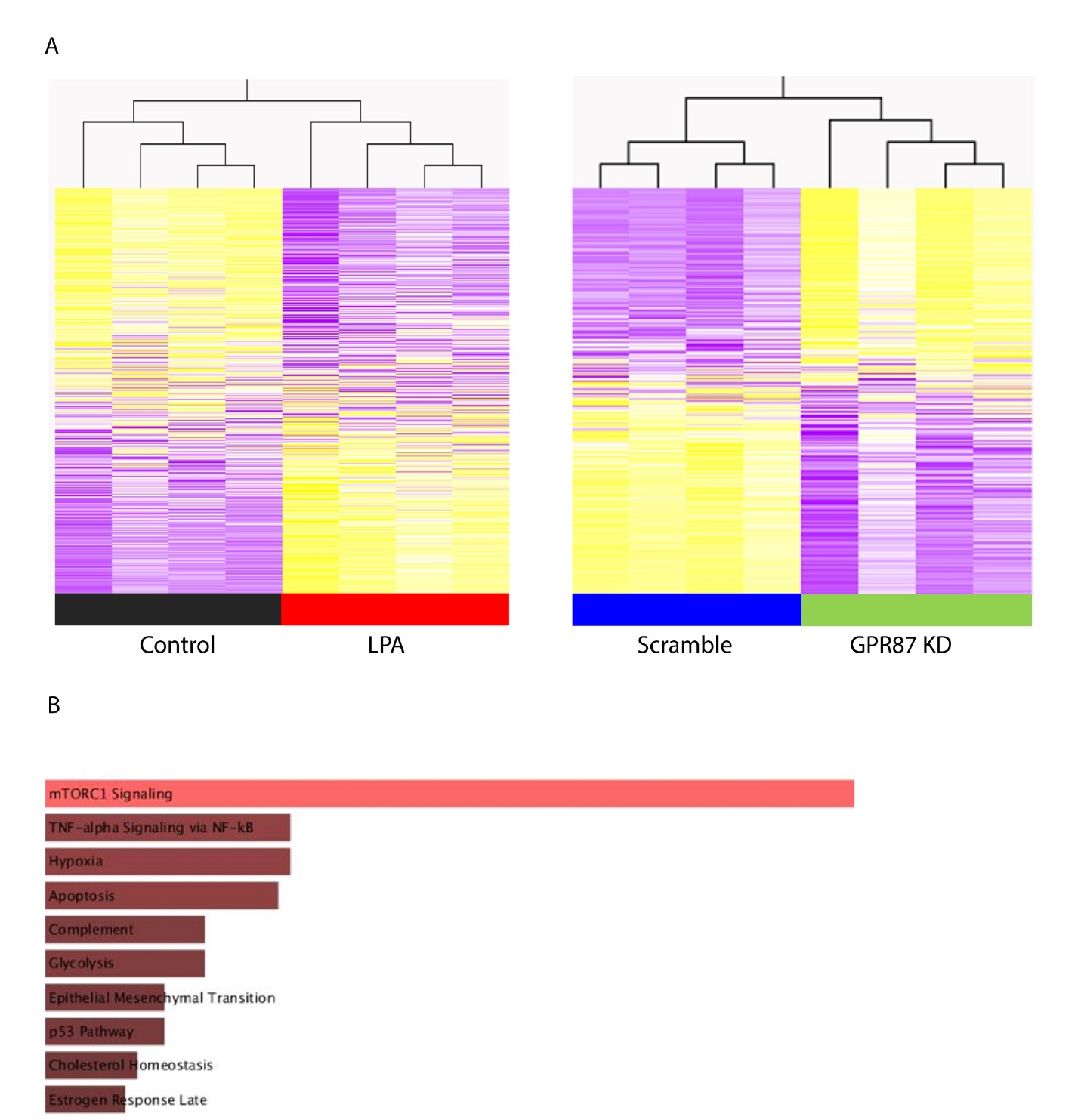

A: B:Top enriched pathways in iBC after GPR87 siRNA knockdown or lysophosphatidic acid treatment.
